# Neural Collective Matrix Factorization for Integrated Analysis of Heterogeneous Biomedical Data

**DOI:** 10.1101/2022.01.20.477057

**Authors:** Ragunathan Mariappan, Aishwarya Jayagopal, Ho Zong Sien, Vaibhav Rajan

## Abstract

**Motivation:** In many biomedical studies, there arises the need to integrate data from multiple directly or indirectly related sources. Collective matrix factorization (CMF) and its variants are models designed to collectively learn from arbitrary collections of matrices. The latent factors learnt are rich integrative representations that can be used in downstream tasks such as clustering or relation prediction with standard machine learning models. Previous CMF-based methods have numerous modeling limitations. They do not adequately capture complex non-linear interactions and do not explicitly model varying sparsity and noise levels in the inputs, and some cannot model inputs with multiple datatypes. These inadequacies limit their use on many biomedical datasets.

**Results:** To address these limitations, we develop Neural Collective Matrix Factorization (NCMF), the first fully neural approach to CMF. We evaluate NCMF on two relation prediction tasks, gene-disease association prediction and adverse drug event prediction, using multiple datasets. In each case, data is obtained from heterogeneous publicly available databases, and used to learn representations to build predictive models. NCMF is found to outperform previous CMF-based methods and state-of-the-art graph embedding methods for representation learning in our experiments. Our experiments illustrate the versatility and efficacy of NCMF for seamless integration of heterogeneous data.

**Availability:** https://github.com/ncmfsrc/ncmf

## 1 Introduction

The complexity and diversity of biomedical data being produced today is unprecedented. Technological advances, e.g., for high-throughput sequencing and drug screening, have enabled rapid generation of data describing the many interconnected and interacting biomedical *entities* (e.g., genes, diseases, drugs). Biomedical data is naturally represented in the form of matrices, e.g., gene expression counts in samples. Entries in a matrix capture dyadic relations, typically as measurements from an experiment, between *entity instances*, along the row and column dimensions. Similarity or interaction matrices, such as protein-protein interactions, capture associations, often across the same entities. Matrix Factorization (MF) is a commonly used unsupervised technique to analyze latent substructures and obtain low-dimensional *entity representations* (or factors) of the row and column entity instances. MF-based techniques (e.g., Principal Components Analysis and Non-negative Matrix Factorization) are used in applications such as identification of mutational signatures in cancer, single cell expression analysis and many more (Stein-O’Brien *et al*., 2018).

Data integration from multiple sources is required in many applications for various reasons: different sources may provide complementary information and one source may compensate for noise, uncertainty or missing values in another. Previous studies have highlighted the merits of utilizing multiple data sources in revealing underlying biological mechanisms and in predictive modeling, e.g., Deng *et al*. (2020); Xu *et al*. (2020); Güvenç Paltun *et al*. (2021). Integrating biomedical data poses several challenges (Žitnik *et al*., 2019). Biological data is often extremely sparse due to experimental measurements being available only on a small subset of exponentially large spaces of possible entity interactions. In many cases, there are a large number of measurements on a limited number of subjects leading to datasets with high dimensions and only few samples. Data is often incomplete, noisy and biased.

Numerous methods have been designed to integrate additional information into MF-based techniques such as those based on Inductive Matrix Completion (Li *et al*., 2020, 2021), Graph-regularized Matrix Factorization (Zhang *et al*., 2020), and Collaborative Matrix Factorization (Wu *et al*., 2020). The typical setting in these methods comprises a central matrix containing known and unknown values of the association of interest (e.g., gene-disease or drug-target) and various ways of incorporating auxiliary data (as features or networks) of the entities of interest (genes and diseases, or drugs and targets) within the modeling framework used. A recent survey can be found in Ou-Yang *et al*. (2021). Most of these methods have been designed for specific contexts and/or datatypes and they cannot be used with *arbitrary* collections of matrices, i.e., collections with any number of input entities and matrices, and where each matrix can have different datatype (see fig. 3 for examples).

The need for integrative analysis with arbitrary collections of matrices arises in many biomedical contexts. For instance, in cancer studies, multi-omics data, including copy number, gene expression, DNA methylation, microRNA and clinical data of several cancers have been collected and analyzed (Weinstein *et al*., 2013) and the integrated analysis of data at different – genomic, epigenomic, transcriptomic and physiological – levels remains a challenge (Senft *et al*., 2017; Xu *et al*., 2019). In such contexts, the requirement is to integrate not only data that is directly related to the entities of interest but also those that are indirectly related. For example, to build a model to identify genes that can be targeted by a drug, one may need to not only use *direct* information about genes and drugs, such as their associations and features, but also *indirect* information such as drug efficacy on patients and clinical data of patients.

Collective matrix factorization (CMF) (Singh and Gordon, 2008) and extensions thereof are models designed to collectively learn from arbitrary collections of matrices. These models generalize the idea of matrix factorization to a collection of matrices. They learn a latent representation for each entity in a way that integrates information from multiple matrices seamlessly. These entity-specific latent representations can be used to perform matrix completion (to predict unknown associations). The latent factors are rich integrative representations that can be used in downstream tasks such as clustering, classification or relation prediction with standard machine learning models, thereby alleviating the resource-intensive task of manual feature engineering from heterogeneous data.

Several extensions of CMF have been developed over the years such as a formulation that enables group-wise sparsity on the latent factors, gCMF (Klami *et al*., 2014), a penalized tri-factorization approach called Data Fusion by Matrix Factorization (DFMF) (Žitnik and Zupan, 2014), and a recent neural approach DCMF (Mariappan and Rajan, 2019). DFMF was empirically evaluated for the task of gene function prediction and DCMF for gene-disease association prediction. In both cases, multiple matrices from heterogeneous sources could be integrated seamlessly for the final prediction task.

However, previous CMF-based methods have numerous modeling inadequacies that limit their application on biomedical datasets, summarized in Table 1. Some of them cannot directly model collections where the same row and column entities are in multiple matrices, a condition we call *Matrix Entity Multiplicity*. For instance, one may want to collectively use two or more Drug-Disease matrices with different data types, representing different associations (e.g., side effects and treatments). DFMF and DCMF do not model mixed (real and categorical) datatypes and most of the previous techniques do not explicitly model noise in the data. None of the previous techniques explicitly model varying levels of sparsity in the matrices that may arise due to missing values. More details are in §2. Biomedical datasets often require modeling all these conditions to obtain effective entity representations from multiple sources.

**Table 1.** Modeling capabilities of previous CMF-based methods and our proposed model NCMF. See text for details.

|  | CMF | gCMF | DFMF | DCMF | NCMF |
| --- | --- | --- | --- | --- | --- |
| Data Sparsity | – | – | – | – | ✓ |
| Noise | – | ✓ | – | – | ✓ |
| Non-linear interactions | partial | – | – | – | ✓ |
| Matrix Entity Multiplicity | ✓ | – | partial | – | ✓ |
| Multiple Datatypes | ✓ | ✓ | – | – | ✓ |

Another significant modeling limitation of previous methods is in their ability to model complex non-linear interactions. The CMF model proposes the use of a link function that is applied after a linear combination of latent factors. In DCMF, non-linearity is modeled with respect to entity representations, but not with respect to matrix factorization – each reconstruction is a linear function (inner product) of the entity representations. DCMF was found to empirically outperform CMF and gCMF. However, none of them can adequately capture complex non-linear dependencies. In other domains, e.g., collaborative filtering (He *et al*., 2017), fully neural methods to generalize matrix factorization have been effective in modeling non-linear interactions. These methods are designed for recommendation systems and do not generalize to arbitrary collections of matrices. To our knowledge, there is no previous fully neural method for Collective Matrix Factorization.

In this paper, we develop Neural Collective Matrix Factorization (NCMF), that is, to our knowledge, the first fully neural approach to CMF that addresses *all* the limitations discussed above. NCMF has a novel architecture that is dynamically constructed based on the number of entities and matrices in the input collection. Through the use of Variational Autoencoders (VAE) where the decoded output is modeled using Zero-Inflated distributions, NCMF effectively models sparse and noisy inputs. Multiple such VAEs are used, one for each row and column for each matrix – this architecture effectively models varying datatypes, sparsity and noise levels across the matrices and matrix entity multiplicity. These VAE representations are fused to form entity-specific representations which are then used to reconstruct the matrix using a neural network, thereby modeling non-linearities in both entity representation learning and matrix reconstruction. The entire network can be trained in an end-to-end manner.

We evaluate NCMF on two important biomedical applications: genedisease association prediction, adverse drug event prediction. In each case, data is obtained from heterogeneous publicly available databases, and the final prediction task is modeled as a relation prediction problem. In terms of predictive accuracy NCMF is found to outperform previous CMF-based methods and state-of-the-art graph embedding methods in our experiments. Clustering and visual inspection of NCMF representations also indicate its superiority over previous CMF methods. Our experiments illustrate the versatality and efficacy of NCMF in seamless integration of data.

## 2 Background and Related Work

### Multiview Learning

In multiview learning, views refer to measurements for the *same* subjects, that differ in source, datatype or modality. Canonical correlation analysis and matrix factorization have been the basis for many methods, including deep learning models such as DCCAE (Wang *et al*., 2015) and CDMF (Wang *et al*., 2017). A recent survey can be found in Nguyen and Wang (2020). These methods cannot model arbitrary collections of matrices.

### Collective Matrix Factorization (CMF)

For a single *m* × *n* dimensional matrix *X*, a low-rank factorization aims to obtain latent factors 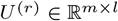, 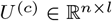, for its row and column entities respectively, such that *X* ≈ *U*^(*r*)^ · *U*^(*c*)*T*^, where *l* < min(*m, n*). This notion is generalized to arbitrary collections of matrices in Collective Matrix Factorization (CMF) (Singh and Gordon, 2008). The *m*^th^ matrix in the collection is factorized as *X*^(*m*)^ ≈ *f_m_* (*U*^[*r_m_*]^ · *U*^[*c_m_*]*T*^) where *r_m_, c_m_* represent the entities along its row and column dimensions respectively and *f_m_* is a matrix-specific link function applied elementwise after multiplying the learnt latent factors. Any two matrices sharing the same entity (e.g., the 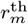 entity) uses the same low-rank representation (*U*^[*r_m_*]^ as part of the approximation, which enables sharing information. This is also called *augmented multiview learning* since it generalizes the multiview setting – auxiliary data from any, even indirectly related, input matrix may be utilized to learn shared representations.

CMF is categorized as an *intermediate integration* fusion technique which learns joint entity-specific representations from multiple input datasets. These representations, in turn, can be used in downstream modeling. In contrast, in early integration, input matrices are fused into a single matrix, e.g., through projection or concatenation, at the data level before subsequent analysis. In late integration, each matrix is used independently for modeling and predictions from the models are combined, e.g., through majority voting. For details, see, e.g., Žitnik *et al*. (2019).

To model view-specific noise and shared structure in some of the matrices, a Bayesian formulation with group-wise sparse priors was designed in gCMF (Klami *et al*., 2014). It can model both Gaussian and non-Gaussian observations, including count and binary data. While gCMF models sparsity at the representation level, sparse inputs are not explicitly modeled. A penalized tri-factorization approach that models matrix-specific associations across entities and constraints that relate objects of the same entity, DFMF was designed by Žitnik and Zupan (2014). Using three factors, i.e., *X*^(*m*)^ ≈ *U*^[*i*]^ · *S_ij_* · *U*^[*j*]*T*^, where *i* = *r_m_, j* = *c_m_*, allows DFMF to model asymmetric relations (e.g., a gene-disease matrix and a different disease-gene matrix) between the same two entities that differ in the third factor (*S_ij_*). However, the general case of multiple matrices with the same row and column entities is not modeled. CMF models such inputs through link functions and Liany *et al*. (2020) develop preprocessing techniques which can be used before applying CMF-based methods to handle such cases.

DCMF, a neural approach to CMF, by Mariappan and Rajan (2019), uses a collection of autoencoders to obtain entity representations from arbitrary collections of matrices. For each entity, all the matrices containing the entity are concatenated (considering transposes wherever required to ensure that the entity remains along the row dimension). DCMF factorizes *X*^(*m*)^ ≈ *g*^[*r_m_*]^(*C*^[*r_m_*]^) · *g*^[*c_m_*]^(*C*^[*c_m_*]*T*^), where *g* is the encoder corresponding to the entity and *C* represents the concatenated matrices containing the corresponding (row *r_m_* or column *c_m_*) entities. The entire network is dynamically constructed – the number of autoencoders is equal to the number of entities in the input – and trained collectively by minimizing the sum of autoencoder reconstruction losses and matrix reconstruction losses for all entities and matrices respectively. Although the entity representation learning is non-linear (through *g*), the matrix reconstruction is alinear function of the representations. Note that in CMF, non-linear interactions may be partly modeled through the link function after the linear inner product. Thus, neither CMF nor DCMF adequately models non-linear interactions.

### Heterogeneous Network Embedding (HNE)

Arbitrary collections of relational data may be represented through Heterogeneous Information Networks (HIN) with multiple types of edges and nodes. Representation learning on HIN aims to obtain vectorial representations of the nodes, called Heterogeneous Network Embeddings (HNE), in a way that preserves the structure and semantics of the HIN.

A detailed survey of HNE methods can be found in Yang *et al*. (2020) who categorize HNE methods into 3 groups: (i) *Proximity-preserving* methods: these are broadly based on random walk approaches (inspired by homogeneous graph embedding methods Deepwalk (Perozzi *et al*., 2014) and LINE (Tang *et al*., 2015)). Metapath2vec (Dong *et al*., 2017) and HIN2vec (Fu *et al*., 2017) use metapath schemes to define semantically relevant walks on HIN from which representations are learnt. Specifying metapaths often require domain knowledge that limits the use of these methods. (ii) *Message-passing* methods: These are typically based on graph neural networks. Examples include Relational Graph Convolutional Network (R-GCN) (Schlichtkrull *et al*., 2018) and Heterogeneous Graph Transformer (HGT) (He *et al*., 2017). (iii) *Relation-learning* methods: these methods have been developed to learn representations of Knowledge Graphs that are essentially HIN. One of the earliest methods, TransE (Bordes *et al*., 2013) designs a method that preserves vectorial translational distance among the edge embedding and embeddings of the incident nodes. This idea has been refined and generalized in many subsequent works, e.g., the use of deep networks in ConvE (Dettmers *et al*., 2018).

A large number of HNE methods have been developed in many different domains and for diverse applications. Any HIN can also be represented as a collection of matrices: each edge type represents a relation (a matrix) and each node type is an entity. Matrices readily accommodate multiple datatypes which requires modeling multiple edge attributes (real-valued weights or categorical labels) in HNE methods. Vectorial node features can easily be added as additional matrices in CMF methods. Both these are particularly relevant for biomedical data that often contain edge information (such as real-valued significance values or ordinal level of an interaction) and node information (auxiliary features about entities).

Among the 3 categories of HNE methods, message-passing methods are more general and most can model node features. Some methods, e.g., R-GCN, can be modified to model weighted edges. However, this often requires changes in their models. Such modifications are less straightforward in proximity-preserving and relation-learning methods. In contrast, our proposed method NCMF can model arbitrary number and types of node and edge features – for example, a new edge feature can be added as another input matrix with no change in the algorithm.

### VAE and DCA

We briefly describe variational autoencoder (VAE) (Kingma and Welling, 2014) and deep count autoencoder (DCA) (Eraslan *et al*., 2019) which we use within the neural architecture of NCMF.

VAE is a generative model that models data *Y* by generating (lower dimensional) latent variables *z*, such that there is a high probability to recover the original data *Y* from *z*. The distribution *P*(*z*|*Y*) is theoretically the best choice to generate *z* but is intractable. So, a variational probability *Q*(*z*|*Y*) is used to approximate the posterior by minimizing the Kullback-Leibler (KL) divergence *D*[*Q*(*z*|*Y*)∥*P*(*z*|*Y*)] which is equivalent to maximizing the variational lower bound log *P*(*Y*) – *D*[*Q*(*z*|*Y*)∥*P*(*z*|*Y*)] given by:

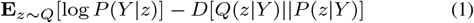

Neural networks are used as probabilistic encoder *Q*(*z*|*Y*) and decoder *P*(*Y*|*z*) and the reparameterization trick (Eraslan *et al*., 2019) is used to obtain samples from *Q*(*z*|*Y*) and ensure differentiability with respect to *z*. More details are in Kingma and Welling (2014). VAEs have been used previously to integrate biomedical data, e.g., by Way and Greene (2018); Simidjievski *et al*. (2019), but they do not model arbitrary collections.

DCA adapts the standard autoencoder architecture (not VAE) to model sparse count data. This is done by defining the reconstruction error as the likelihood of the distribution for a noise model instead of reconstructing the input data itself. The DCA architecture has a neural network as an encoder and decoder wherein the decoder outputs are the parameters of the noise model instead of the input to the encoder. Neural networks capture intrinsic dependencies, allow low-dimensional reprsentation learning and the noise model allows unsupervised learning of noise and signal in the data. Eraslan *et al*. (2019) recommend the use of the Zero Inflated Negative Binomial (ZINB) distribution for ordinal data. We provide details for ZINB here, and Zero Inflated Normal (ZIN) distribution, used for real-valued data, is described in Appendix 1.

ZINB is a mixture of 2 components: (i) a point mass at zero representing the zero values and (ii) a negative binomial component for ordinal values. The ZINB distribution is parameterized by the mean (Ω) and dispersion (Θ) of the negative binomial (NB) and the mixture coefficient (Π) that represents weight of the point mass: 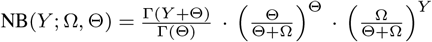 where Γ is the gamma function. ZINB(*Y*; Π, Ω, Θ) = (Π · Δ_0_(*Y*)) + ((1 – Π) · NB(Y; Ω, Θ)) where Δ_0_ is the point mass distribution at 0. Thus, ZINB models *Y* = 0 with probability Π and *Y* ~ *NB* with probability 1 – Π.

## 3 Neural Collective Matrix Factorization (NCMF)

The inputs to NCMF are (i) an arbitrary collection of *M* matrices containing relational data among *N* entities and (ii) an entity-matrix bipartite relationship graph 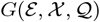, where the vertices 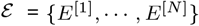 represent entities, 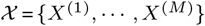 represent matrices and edges 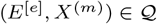 show the row and column entities ineachmatrix. Fig. 2 shows an example–a collection with *M* = 5, *N* = 6 and its entity-matrix relationship graph (*E*^[1]^ is shaded to show the corresponding node). Each entity has multiple *instances* corresponding to rows or columns in one or more matrices. We use *e* to index entities and *r_m_, c_m_* represent the row and column entities, respectively, in the *m*^th^ matrix (*e, r_m_, c_m_* ∈ {1,…, *N*}). We use [*e*], (*m*) and (*e, m*) to denote entity *E*^[*e*]^, matrix *X*^(*m*)^ and edge (*E*^[*e*]^, *X*^(*m*)^) respectively. Let *d*[_*e*_] represent the number of instances of the *e*^th^ entity. NCMF collectively learns entity representations from the inputs and performs matrix completion to obtain:

1. 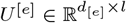: Latent l-dimensional representations of all instances of the *e*^th^ entity.
2. 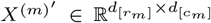: Reconstructed matrix corresponding to each of the matrices *X*^(*m*)^.

### 3.1 Dynamic Network Architecture

To enable learning from *arbitrary* collections, the NCMF network is dynamically constructed based on the inputs as shown in fig. 1 and described in the following. There are 3 subnetworks in our architecture: (i) Autoencoder Subnetwork: 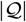 autoencoders to obtain row-wise and column-wise entity representations in each matrix; (ii) Fusion Subnetwork: 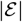 feedforward networks to fuse multiple encodings of entities; (iii) Matrix Completion Subnetwork: 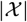 feedforward networks to reconstruct the input matrices. Note that 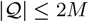, 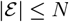, 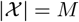.

**Fig. 1:**
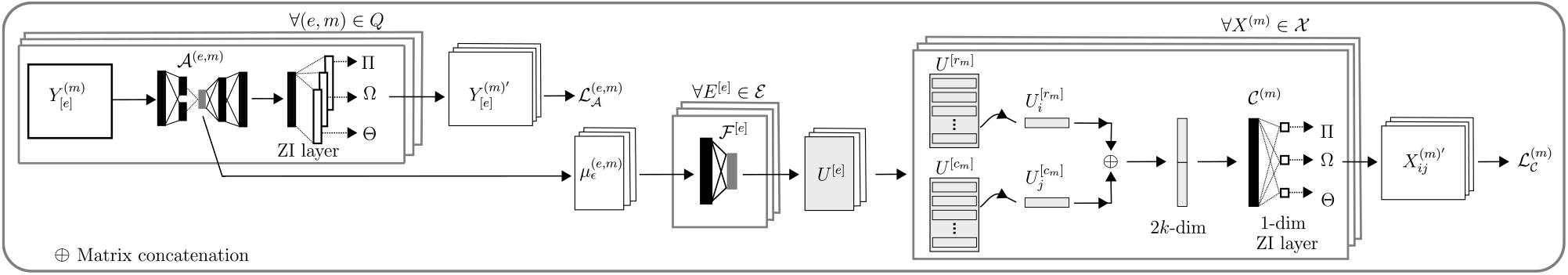
Schematic of NCMF Network Construction and Training

**Fig. 2:**
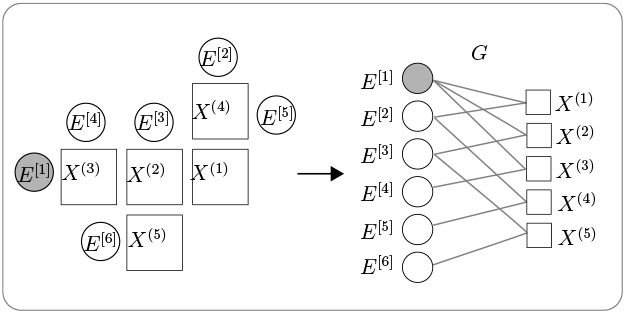
Collection with 6 entities, 5 matrices and its entity-matrix relationship graph G.

#### Autoencoder Subnetwork

This subnetwork, denoted by 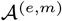, consists of autoencoders, one for each entity in each matrix. In the *m*^th^ matrix, the row vectors are inputs to the row-entity autoencoder and the column vectors are the inputs to the column-entity autoencoder. The same entity may be present in multiple matrices obtained from diverse sources which may differ in datatypes, noise and sparsity levels. So, we model them separately using different autoencoders. The same entity may also be present within a matrix, e.g., in adjacency matrices from graphs. By obtaining row and column entity representations separately we capture correlations across both dimensions.

Let 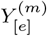 denote the *e*^th^ entity instances (in rows or columns) in the matrix *X*^(*m*)^ along its rows. For each autoencoder in 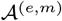, we use the underlying probabilistic framework of VAEs to model complex, high-dimensional data and adapt it for sparse, noisy inputs. We assume a Gaussian latent representation 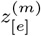 for *e*^th^ entity of the *m*^th^ matrix.

We use a neural network *f_ϵ_* as a probabilistic encoder 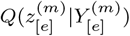 and infer its mean and standard deviation. For the probabilistic decoder, 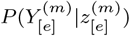, we use Zero-Inflated distributions to model noisy, sparse and overdispersed input data. We use a neural network *f_δ_* as a probabilistic decoder to obtain 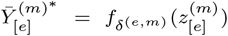. This is used as input simultaneously to the final layer with 3 outputs to learn the 3 parameters of ZINB. The mean and dispersion parameters are always non-negative, so we use an exponential activation function. To ensure that the weight Π remains between zero and one, sigmoid activation is used.

- 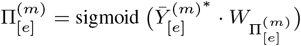
- 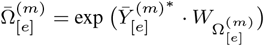
- 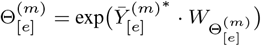

The encoder can be viewed as a parameterized link function specific to an entity for each matrix. The latent variable mean 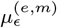 is considered to be the representation of the *e*^th^ entity in the *m*^th^ matrix.

#### Fusion Subnetwork

For the final matrix reconstruction, we require a single entity representation that is obtained by the fusion subnetwork denoted by 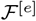. If the same entity is present in more than one matrix, the final entity-specific representation is obtained by 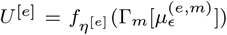, where Γ*_m_*[.] represents the concatenation operation and the index *m* iterates over all matrices containing the *e*^th^ entity. If the entity is present in a single matrix, no fusion is required and we set 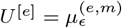.

#### Matrix Completion Subnetwork

This subnetwork, denoted by 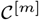, has one network for each matrix in the input. Each matrix *X*^[*m*]^ is reconstructed using its row and column entity representations, *U*^[*r_m_*]^, *U*^[*c_m_*]^. To learn non-linear interactions between them, we use another feedforward network, *f_γ_*, for the *ij*^th^ matrix entry: 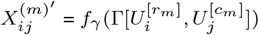 where Γ[.] represents the concatenation operator applied on the *i*^th^ row and *j*^th^ column entity representations. To model the noise and sparsity in the matrix, we add a final layer to learn three 1-dimensional ZINB parameters (or ZIN, in case of real-valued matrices) for each matrix entry, Weight (Π), Dispersion(Θ) and Mean(Ω). As described earlier, an exponential activation is used for mean and dispersion outputs and a sigmoid activation is used for the weight output. The reconstructed entry is the mean output from the final layer.

### 3.2 Network Training

The loss function to be minimized is 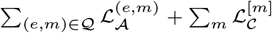. The first term sums the autoencoder losses in subnetwork 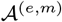 and the second term sums the matrix reconstruction losses in subnetwork 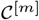.

Our 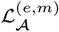 corresponds to the negative of the variational lower bound given by Eq. 1. By assuming a Gaussian 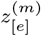, the expression for 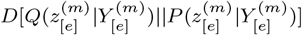 corresponding to Eq. 1 follows directly from those in Kingma and Welling (2014). A Monte Carlo estimate of the expectation in Eq. 1 can be obtained through the ZINB likelihood for a minibatch of samples. This is interpretable as the expected negative reconstruction error with the KL divergence term acting as a regularizer. Empirically we find that adding another ridge regularizer on the weight parameter improves performance, as suggested by Eraslan *et al*. (2019). Thus, we have 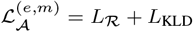,

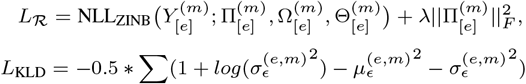

where NLL_ZINB_ is the negative log likelihood of ZINB distribution and *L*_KLD_ is the KL Divergence term. The matrix reconstruction loss is also calculated using the ZINB likelihood and the ridge regularizer: 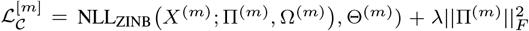. In case of real inputs, ZIN likelihoods are used instead of ZINB.

#### Algorithm 1: Neural Collective Matrix Factorization (NCMF)

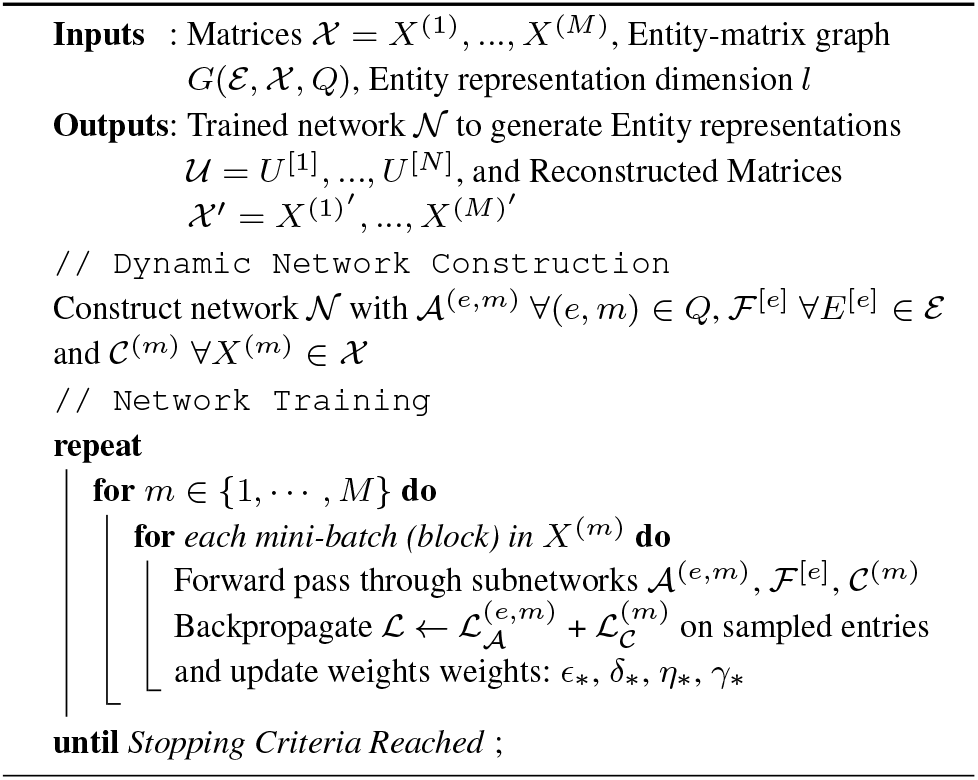

For each matrix in the collection, we create mini-batches by considering a *block* within the matrix. A *block* is a set of row and column entity instances along with the corresponding matrix entries. All the matrix rows and columns that participate in the block are considered in each minibatch, i.e., if the *ij*^th^ entry of the matrix is contained in a block then the entire *i*^th^ row and *j*^th^ column is considered in the mini-batch. The entire row (column) is used as input in the autoencoder corresponding to the row (column) of the matrix, in the autoencoder subnetwork. Only the entries in the block are reconstructed in the matrix completion subnetwork, using the representations of the corresponding *i*^th^ row and *j*^th^ column entity instances. During loss computation, we found that subsampling the zero entries led to improved performance. Within each block, if there are *n* non-zero entries, we sample nk zero entries and use only these *n*(1 + *k*) entries for computing 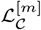. The loss 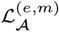 is computed for the rows and columns used in the autoencoders.

We randomly initialize model parameters and optimize the model with AdamW (Loshchilov and Hutter, 2017a). We use cosine annealing with warm restarts for the learning rate (Loshchilov and Hutter, 2017b) and gradient norm clipped to 1. Training terminates when the difference in loss in consecutive iterations is less than a fixed threshold or if the maximum number of epochs is complete. Algorithm 1 has an outline of NCMF.

NCMF improves over the neural architecture of DCMF and overcomes several limitations of previous methods (summarized in Table 1). NCMF explicitly models noise and sparsity in input data through VAEs and Zero Inflated distributions. Through a fully neural architecture, and, in particular, neural layers for generalized matrix factorization, NCMF models non-linearity both at the entity representation and matrix factorization levels. The neural architecture also allows arbitrary number of input matrices (including those with the same row and column entities), and supports multiple input data types.

## 4 Case Studies and Data

We evaluate NCMF on two biomedical applications: gene-disease association prediction, adverse drug event (ADE) prediction where we consider side-effects of both single and multiple drugs (aka polypharmacy). In each case, data is obtained from heterogeneous databases and the task is modeled as a relation prediction problem. A schematic of the constituent matrices and entities in the data is in fig. 3. that also illustrates the versatality of NCMF in terms of its ability to learn from large number of matrices (PubMed data, 10 matrices), collections with different kinds of relations among the same number of matrices (MIMIC and PolyP datasets).

**Fig. 3:**
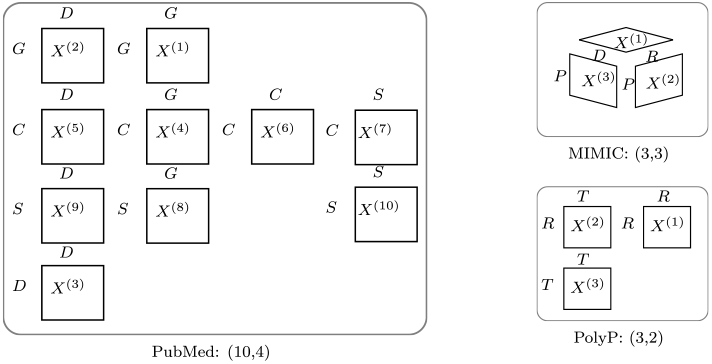
Datasets; (*M, N*): number of matrices and entities.

### 4.1 Gene-Disease Association Prediction

Knowledge of genes associated with diseases not only improves our understanding genomic interactions but also facilitates the design of treatment strategies. Many experimental methods have been developed to determine such associations but they are expensive, time-consuming and may be specific to certain classes of diseases (Piro and Di Cunto, 2012). Hence, computational approaches are used to first prioritize the candidate pairs before experimental verification, including matrix completion based methods that can use heterogeneous auxiliary data sources, e.g., Natarajan and Dhillon (2014); Mariappan and Rajan (2019).

We use the benchmark *PubMed* dataset comprising relations across 4 entities – genes, diseases, chemicals and species. Both the entities and their relations have been inferred from biomedical literature, by Natural Language Processing on articles from PubMed (Canese and Weis, 2013). Named entity recognition and relation pattern mining methods were used in combination with manual curation to create the final dataset. More details can be found in Yang *et al*. (2020). Fig. 3 shows a schematic of the input matrix collection. We randomly sample 3 datasets from the collection, each with the dimensions listed in Table 2. Sparsity levels (percentage of zeros) are for one of the datasets, and are very similar in the other two.

**Table 2:** PubMed Dataset Statistics (G: Gene, D: Disease, C: Chemical, S: Species, Col: Column, Dim: Dimension)

| Matrix | Row | Col | Row Dim | Col Dim | Sparsity% |
| --- | --- | --- | --- | --- | --- |
| $X^{(1)}$ | G | G | 2661 | 2661 | 99.94 |
| $X^{(2)*}$ | G | D | 2661 | 4288 | 99.96 |
| $X^{(3)}$ | D | D | 4288 | 4288 | 99.97 |
| $X^{(4)}$ | C | G | 5546 | 2661 | 99.97 |
| $X^{(5)}$ | C | D | 5546 | 4288 | 99.97 |
| $X^{(6)}$ | C | C | 5546 | 5546 | 99.98 |
| $X^{(7)}$ | C | S | 5546 | 592 | 99.98 |
| $X^{(8)}$ | S | G | 592 | 2661 | 99.97 |
| $X^{(9)}$ | S | D | 592 | 4288 | 99.97 |
| $X^{(10)}$ | S | S | 592 | 592 | 99.98 |

### 4.2 Adverse Drug Event (ADE) Prediction

ADEs are unintended drug side effects that often lead to emergency visits, prolonged hospital stays, and worse patient outcomes (Ventola, 2018). They pose tremendous clinical and economic burden – in the US alone morbidity and mortality costs associated with ADEs are estimated to be approximately $528 billion in 2016 (Watanabe *et al*., 2018). Clinical trials are limited by the number and characteristics of patients tested, the duration of the observation period and may not detect all ADEs, especially those with long latency or those that affect only certain patient groups (Coloma *et al*., 2013). Therefore, post-marketing drug safety surveillance is routinely conducted to detect unknown ADEs from Electronic Medical Records (EMR) and other sources. We consider two different data sets for single drug and polypharmacy side-effect predictions.

#### Drug Side Effect Prediction from EMR Data

We use MIMIC-III, a freely accessible EMR database of patients admitted to Intensive Care Units (Johnson *et al*., 2016). Two structured tables – diagnoses and prescriptions – are used from MIMIC, that contain the diagnosed diseases and prescribed drugs respectively. ADE relations are obtained from another database, OFFSIDES (Tatonetti *et al*., 2012), that contains off-label side effects collected from doctors, patients and drug companies in adverse event reporting systems. The two databases follow different coding systems that had to be mapped for us to use them collectively. MIMIC follows normalized NDC and short ICD9 codes while OFFSIDES follows RxNORM and decimal ICD9 codes for drugs and diseases respectively. We used the mapping provided by Kury and Bodenreider (2017) to map NDC to their RxNORM codes. We converted the decimal ICD9 codes in OFFSIDES to short ICD9 using the icd9 library (Shedden, 2015). After mapping, we found 596 unique drugs and 1321 unique diseases common across MIMIC and OFFSIDES.

There were 34,807 patients who were administered these 596 drugs and had 1-5 drugs with at least 5 or more side effects listed in OFFSIDES. Since ADEs are associated with longer Length of Stay (LoS) (Davies *et al*., 2009), we created a dataset containing patients with both low and high LoS. The mean LoS of all admissions was 258 hours. We sampled three groups of patients, one with LoS less than or equal to 258, another with LoS greater than or equal to 500 (mean + 1 standard deviation) and a third group with LoS between 258 and 500 hours. These random selections were done three times to obtain 3 sampled datasets. A patient may have multiple admissions and we randomly selected one admission per patient. The dataset statistics for one of the samples are shown in Table 3.

**Table 3:**
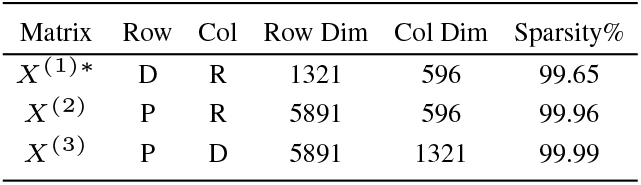
MIMIC Data (P:Patient, D: Disease, R: Drug)

#### Polypharmacy Side Effect Prediction

Patients may be prescribed multiple drugs together, when they have coexisting ailments or for combination therapies, e.g., in cancer (Mokhtari *et al*., 2017). In such cases, polypharmacy ADEs may occur due to due to drug-drug interactions (Tatonetti *et al*., 2012).

We use the benchmark polypharmacy dataset of Žitnik *et al*. (2018) consisting of three types of interactions: drug-drug, drug-protein and protein-protein interactions. Drug-drug interactions contain triplets of the form drugA-SE-drugB where consuming drugA and drugB together would cause the side effect (SE) mentioned in the triplet. E.g., aspirin - Kidney-Failure - warfarin. This data has been curated from multiple databases: SIDER (Kuhn *et al*., 2016) and TWOSIDES (Tatonetti *et al*., 2012). Drug-protein interactions contain experimentally verified small chemicals (drugs) that target specific proteins. Protein-protein interactions are curated from multiple databases indicate physical interactions experimentally documented in humans. More details can be found in Žitnik *et al*. (2018).

We create two datasets as follows. In the first dataset, that we call *PolyP1*, we use the preprocessed benchmark dataset, including the exact train-test splits provided by Burkhardt *et al*. (2019) and used in Dasgupta *et al*. (2021), the methods that, to our knowledge, have the best results on this data. This data considers only the most common among the 963 side effects present in at least 500 drug combinations. This resulted in 645 drugs and their associated side effects. There are 22,583 proteins. Drug-protein and protein-protein matrices are binary, indicating presence or absence of interactions. The drug-drug matrix can have 964 values, indicating no side-effect or one of the 963 side-effects. For each drug-drug pair, the prediction task is predict which among the 963 side effects, or the absence of any side-effect, would occur.

Some baseline methods that we compare with cannot learn sideeffect representations from the drug-drug matrix, for use in downstream prediction tasks. To have a dataset to compare all the methods on, we extract a second dataset, that we call *PolyP2* in the following manner. The drug-drug matrix is binarized, i.e., for a drug-drug pair, presence of a side effect is indicated by 1 and absence by 0 and the task is that of predicting whether or not a polypharmacy side effect occurs. We use the same 645 drugs and only those proteins that interact with more than 20 drugs are used, resulting in 837 proteins. Datasets statistics are in Table 4.

**Table 4:** PolyP2 Data Statistics (R: Drug, T: Protein) ^*^: matrix used for evaluation

| Matrix | Row | Col | Row Dim | Col Dim | Sparsity% |
| --- | --- | --- | --- | --- | --- |
| $X^{(1)*}$ | R | R | 645 | 645 | 75.53 |
| $X^{(2)}$ | R | T | 645 | 837 | 97.11 |
| $X^{(3)}$ | T | T | 837 | 837 | 97.10 |

## 5 Experiment Settings

We compare the quality of the representations, obtained by NCMF to those from previous CMF-based and state-of-the-art HNE methods. All hyperparameter settings used in our experiments are in Appendix 2.1.

### Comparison with CMF Methods

We compare the performance of NCMF with 4 previous CMF methods: CMF (Singh and Gordon, 2008), gCMF (Klami *et al*., 2014), DFMF (Žitnik and Zupan, 2014) and DCMF (Mariappan and Rajan, 2019).

#### Relation Prediction

We first compare the performance on relation prediction where the relations in our datasets PubMed, MIMIC and PolyP2 correspond, respectively, to associations of gene-disease, drug-ADE and polypharmacy-ADE. In each case, one of the input matrices is used for evaluation as indicated in Tables 2, 3 and 4. On this matrix, 20% of the relations (both positive and negative) are randomly selected and held out. The remaining 80% of the relations are used along with all other input matrices to obtain entity representations, using each of the CMF-based methods. We call the latter the RL split used for unsupervised *representation learning* and the former EV split used for *evaluating* the representations in a supervised setting as described as follows. During RL-EV split creation, we ensure that all entity instances in the EV set are also present in the RL set. This is done to have representations, which will be used as features, for all entity instances in the EV set. We evaluate 3 such randomly sampled splits.

The representations learnt on the RL split are evaluated following procedure of Yang *et al*. (2020) to evaluate embeddings. For the *uv*^th^ hidden cell, we combine the representations of the *u*^th^ row entity instance and the *v*^th^ column entity instance using the Hadamard function to form a feature vector. The cells in each of the 3 EV sets are divided into five cross validation (CV) folds and in each CV iteration, the constructed features are used to train a linear SVM on four training folds and predict on the fifth fold. We report the 5-fold CV performance of all the compared algorithms averaged over the 3 EV sets. AUC (area under the ROC curve) and MRR (mean reciprocal rank) are used as evaluation metrics.

#### Visualization and Clustering

Second, we visually inspect the representations learnt through NCMF and that of the best-performing baseline CMF method, for each dataset. We use hypertools (Heusser *et al*., 2017) to view the representations in 2 dimensions, using the default option of Incremental PCA to obtain the plots. We also evaluate how well the learnt representations discern intrinsic clusters in the data. Since the representations are high-dimensional, we choose the first five principal components and use them to find clusters using K-Means. In the PubMed dataset, we choose the disease entity for which labels, indicating disease categories, are provided for 241 disease instances by Yang *et al*. (2020). In the MIMIC dataset, we choose the patient entity labelled by the 3 categories of Length of Stay (LoS) as described in $4. In the Polypharmacy dataset, we choose the drug entity. There is no single label provided in the data to categorize drugs. So, we consider the top 5 most frequently associated target proteins as (distinct) binary labels and report the mean performance. With respect to each of these labels, we compute the Adjusted Rand Index (ARI) to evaluate clustering performance.

#### Runtime

Third, we compare the runtime of the methods on each of the datasets. Note that only the RL splits are used for representation learning.

### Comparison with HNE Methods

We first compare the performance of NCMF on the PubMed and PolyP1 benchmark datasets on which the current best performance have been reported by specific HNE methods. To our knowledge, the best performing previous method on the PubMed dataset is ComplEx (Trouillon *et al*., 2016), in (Yang *et al*., 2020), where the same metrics AUC and MRR are used. On the polypharmacy data (PolyP1), the best results are that of Weighted Deepwalk (Dasgupta *et al*., 2021), on the same train-test split used by the best previous methods ESP (Burkhardt *et al*., 2019). On this data, the evaluation metrics used were: AUC, area under the precision-recall curve (AUPRC), and average precision at 50 (AP50) for each of the 963 side effects, which were then averaged. Details of the preprocessing done to apply NCMF are given in Appendix 2.2.

We also compare the performance on relation prediction on PubMed, MIMIC and PolyP1 datasets with 6 HNE methods from all three categories: (i) Proximity-preserving methods: Metapath2Vec (Dong *et al*., 2017), HIN2Vec (Fu *et al*., 2017); (ii) Relation learning methods: ConvE (Dettmers *et al*., 2018), TransE (Bordes *et al*., 2013); (iii) Message passing methods: R-GCN (Schlichtkrull *et al*., 2018), HGT (Hu *et al*., 2020). The experiment setting is identical to that for evaluating CMF-based methods. While we compare the performance of NCMF with several representative HNE methods and state-of-the-art HNE methods for the benchmark datasets, a comprehensive theoretical and empirical comparison with the large number of HNE methods in the literature is beyond the scope of this work and may be addressed in future work.

### NCMF Model Analysis

We evaluate the importance of the neural components within the NCMF architecture through an ablation study. With the same experiment setting used to evaluate CMF-based methods for relation prediction, we evaluate the performance on the PubMed dataset, after making the following changes (separately) to NCMF:

- NCMF-ZI: We remove the Zero-Inflated layers in the autoencoder and matrix completion subnetworks and use RMSE loss instead of the negative loglikelihood in both cases.
- 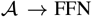: We remove the VAE and the corresponding loss during training. Instead, we use a feedforward network whose output is passed to the fusion subnetwork.
- 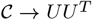: We remove the neural matrix completion subnetwork. For each cell of the matrix, corresponding row and column representations from the fusion layer are used to complete the matrix by multiplication and the RMSE loss is backpropagated instead of the negative loglikelihood.

In each case, the rest of the NCMF network remains the same.

## 6 Results and Discussion

### 6.1 Comparison with CMF methods

Fig. 5 shows the performance of NCMF and baseline CMF methods on all 3 datasets. On both metrics NCMF significantly outperforms other methods (p-value < 0.05, Friedman-Nemenyi test (Demšar, 2006)). DFMF is the second best performing method in the PubMed and MIMIC datasets. DCMF, the other neural approach is the next best in the PolyP2 dataset and is comparable to that of DFMF in the other two.

Fig. 4 shows the 2-dimensional visualizations (obtained via PCA) of disease representations in the PubMed dataset, patient representations in the MIMIC dataset and drug representations in the PolyP2 dataset, where colors depict corresponding cluster labels. The representations from NCMF are compared with those from the next best performing method (with respect to relation prediction). In all cases, representations from NCMF appear to be well spread out. ARI values shown in Table 5 indicate better clustering in two datasets. These results indicate that the representations learnt from heterogeneous inputs capture correlations with respect to the chosen labels well.

**Fig. 4:**
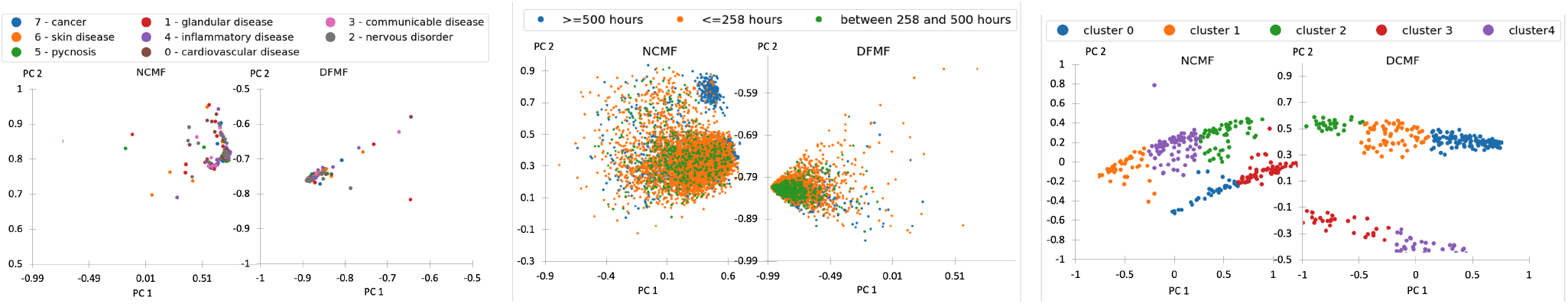
Disease (left), patient (middle) and drug (right) entity representations learnt by NCMF on PubMed, MIMIC and PolyP2 datasets respectively; PC: Principal Component.

**Fig. 5:**
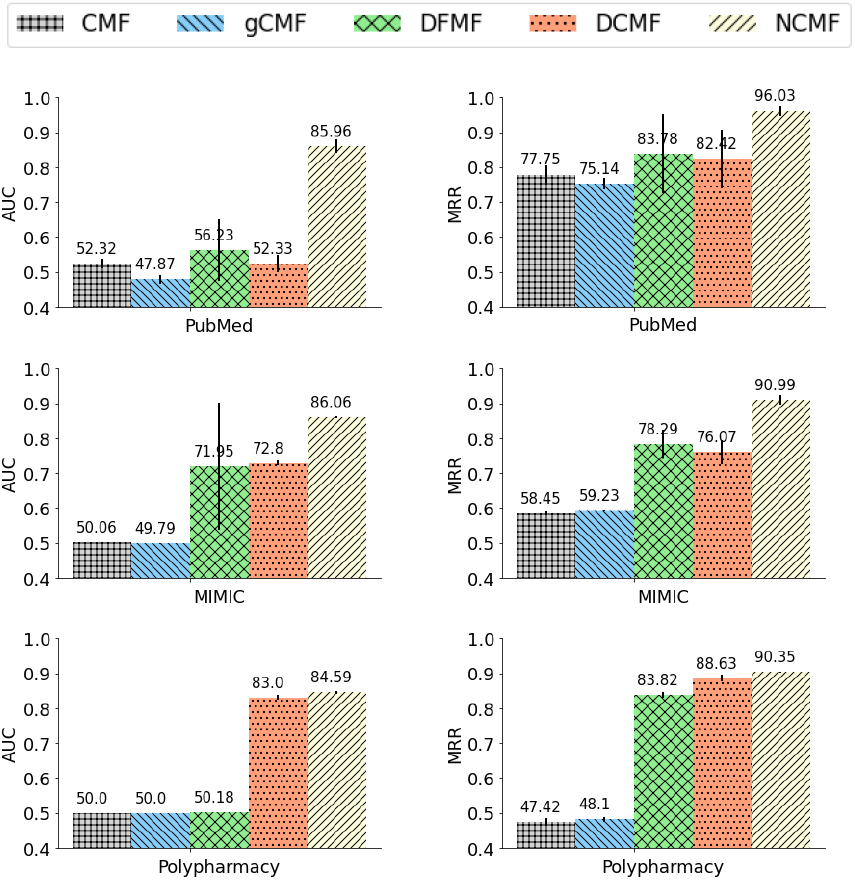
Relation Prediction: Comparison with CMF-based methods.

**Table 5.** ARI values for clustering with NCMF and next-best performing CMF method.

| PubMed |  | MIMIC |  | PolyP2 |  |
| --- | --- | --- | --- | --- | --- |
| NCMF | DFMF | NCMF | DFMF | NCMF | DCMF |
| **0.0015** | 0.0003 | **0.1535** | 0.1063 | 0.1396 | **0.1945** |

We present a detailed analysis of runtime in Appendix 3.1. To summarize, among the CMF-based methods, DFMF is the fastest; the neural methods NCMF and DCMF are comparable in speed and slower than DFMF. Both DCMF and NCMF are faster than CMF and gCMF.

### 6.2 Comparison with HNE Methods

Table 6 shows the performance of NCMF compared to that of the previous best reported methods on these benchmark datasets. On the PubMed dataset, NCMF outperforms ComplEx in both AUC and MRR. On the PolyP1 dataset, NCMF has better average AUC and AUPRC compared to that of Weighted DeepWalk, while the average precision is lower. Appendix 3.3 has additional analysis of predictions from NCMF on the PolyP1 dataset.

**Table 6.**
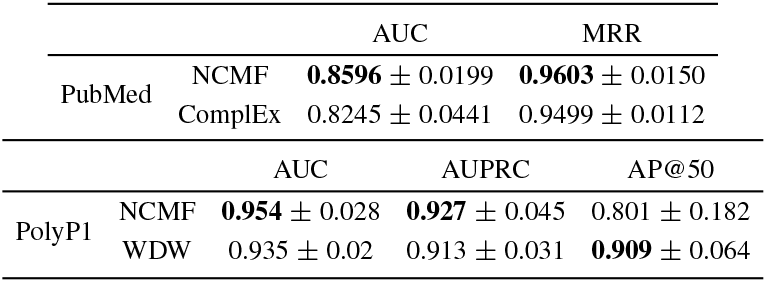
Performance of NCMF on benchmark datasets, compared with the best previous methods. WDW: Weighted Deepwalk.

Fig. 6 shows the performance of NCMF and six HNE methods. NCMF outperforms all six HNE methods in MRR on all datasets and in AUC on 2 out of 3 datasets. Among the HNE methods, HGT and ConvE have the best performance, that are in most cases better than all the CMF-based methods, except NCMF.

**Fig. 6:**
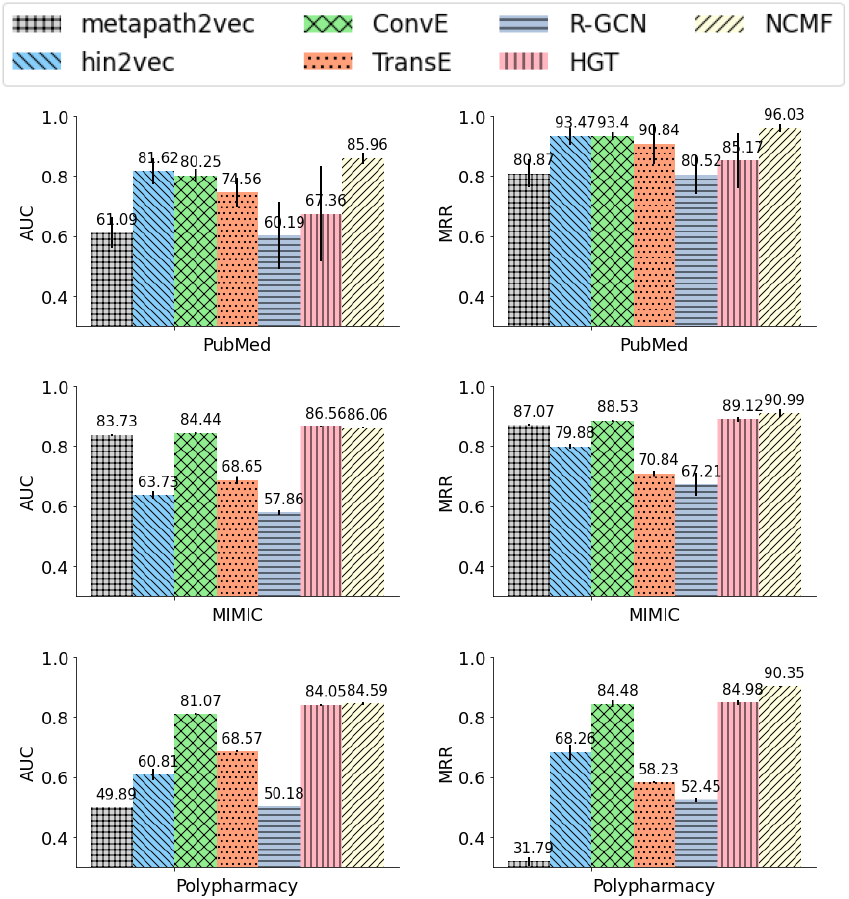
Relation Prediction: Comparison with HNE methods

### 6.3 NCMF Model Analysis

Table 7 shows the results of our ablation study. We observe that in each case, the performance of the modified model is inferior to that of NCMF. Without the ZI layers, the model does not capture noise and sparsity well. Replacing the VAE with a feedforward network essentially removes the decoder and makes the network simpler. However, the decoder is known to have a regularizing effect and its removal deteriorates performance. Without the neural matrix completion layer, a linear model for matrix completion is assumed that does not adequately model non-linear interactions. Overall, these results strongly suggest that every architectural component in NCMF plays an important role in effective predictive modeling.

**Table 7.** Ablation Study: Performance – mean (standard deviation) – on PubMed Dataset.

| | NCMF | NCMF-ZI | $\mathcal{A} \rightarrow \text{FFN}$ | $\mathcal{C} \rightarrow \text{UU}^T$ |
| --- | --- | --- | --- | --- |
| AUC | **0.8596**<br>(0.0199) | 0.6126<br>(0.0418) | 0.7122<br>(0.0178) | 0.8371<br>(0.0351) |
| MRR | **0.9603**<br>(0.0150) | 0.9058<br>(0.0318) | 0.9521<br>(0.0183) | 0.9554<br>(0.0226) |

## 7 Conclusion

We present NCMF, that to our knowledge is the first fully neural method for Collective Matrix Factorization (CMF). NCMF addresses the limitations of previous CMF-based methods in learning representations from matrices varying in sparsity, data types and noise levels. Unlike previous neural CMF methods both representation learning and matrix completion in NCMF are fully neural, enabling better modeling of nonlinear associations. Thus, NCMF is particularly useful in integrative modeling of biomedical data with arbitrary auxiliary information, as seen in our experiments. We evaluate the performance of NCMF on the relation prediction tasks using multiple real datasets curated from heterogeneous sources. NCMF is found to outperform previous CMF-based methods and several representative HNE methods. On the benchmark datasets for polypharmacy side effect prediction and gene-disease prediction, NCMF outperforms previous methods with best known results.

The main limitation of NCMF is its high running time, both for training and hyperparameter tuning, especially on large input collections. This may be partly alleviated through the use of high-performance hardware. More scalable optimization methods and effective hyperparameter tuning techniques may be developed in the future to improve its running time.

## Appendix

### 1 Zero Inflated Normal Distribution

ZIN is a mixture of 2 components: (i) a point mass at zero representing the zero values and (ii) a Normal component for real values. The ZIN distribution is parameterized by the mean (Ω) and dispersion (Θ) of the Normal (N) and the mixture coefficient (Π) that represents weight of the point mass: 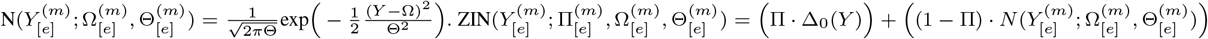 where Δ_0_ is the point mass distribution at 0. Thus, ZIN models *Y* = 0 with probability Π and *Y* ~ *N* with probability 1 – Π.

### 2 Code, Data and Experiment settings

#### 2.1 NCMF Experiment Settings

The code, data, and instructions needed to reproduce all experimental results are in the repository https://github.com/ncmfsrc/ncmf. Table 8 lists the hyperparameters used for NCMF in our experiments for each dataset. These were chosen through hyperparameter tuning, for which the ranges and choices made are listed in Table 9.

**Table 8.** Final Hyperparmeters used to run NCMF with the datasets

| Dataset → | PubMed | MIMIC | PolyPharmacy |
| --- | --- | --- | --- |
| **Hyperparameter ↓** |  |  |  |
| Learning rate | 1e-5 | 1e-6 | 1e-4 |
| Weight Decay | 1e-4 | 5e-1 | 1e-4 |
| Batch Size | 1024 | 2048 | 1024 |
| Number of Iterations | 350 | 1000 | 1000 |
| Number of encoding and decoding layers | 3 | 3 | 3 |
| Encoding/Decoding activation function | tanh | tanh | relu |
| Entity representation size | 100 | 50 | 50 |
| Convergence criterion | $-1e^{-3}$ | $-1e^{-3}$ | $1e^{-6}$ |

**Table 9.** Range of hyperparmeters tried with NCMF by trial and error, before manually selecting the final based on the validation set’s RMSE of the matrix reconstruction

| Dataset → | PubMed | MIMIC | PolyPharmacy |
| --- | --- | --- | --- |
| **Hyperparameter ↓** |  |  |  |
| Learning rate | 1e-6 to 1e-3 | 1e-6 to 1e-3 | 1e-6 to 1e-3 |
| Weight Decay | 1e-4 to 0.5 | 1e-4 to 0.5 | 1e-4 to 0.5 |
| Batch Size | 1000 to 2800 | 1000 to 2800 | 1000 to 2800 |
| Number of Iterations | 100 to 1000 | 100 to 1000 | 100 to 1000 |
| Number of encoding and decoding layers | 3 | 3 | 3 |
| Encoding/Decoding activation function | relu,tanh | relu,tanh | relu,tanh |
| Entity representation size | 50,100 | 50,100 | 50,100 |

All the experiments were run using Intel(R) Xeon(R) Gold 6130 CPU @ 2.10GHz and NVIDIA Tesla V100S PCIe 32 GB graphics card. To enable replication of results, we used random seeds and are hardcoded in the source files. Table 10 lists the methods (where applicable) and the corresponding random seed values.

**Table 10.**
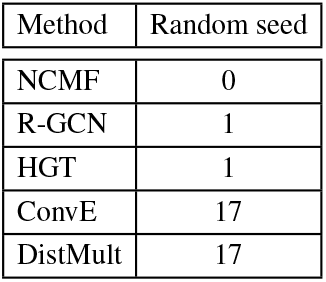
Random seed values

#### 2.2 NCMF on Polypharmacy Data

The protein-protein and drug-protein matrices, of dimensions 22583 × 22583 and 645 × 22583 respectively, are used directly from the PolyP1 dataset. The *ij*^th^ cell in the drug-drug matrix indicates the side effect associated with the pair (drug i, drug j). The cells can have 964 possible values, indicating 963 possible side effects or no side effect. The data is extremely sparse and there is insufficient signal to discern all 964 cases through matrix completion alone. Also note that the three matrices have just two entities – drug and protein. Using just the drug representations (from CMF or HNE methods) as features, we cannot use side effects as labels in a multi-label classification model, since the same drug pair can have multiple side effects.

To use NCMF, we create a new entity called drugSE. Each drugSE instance represents a pair (drug, side effect) and there are 645 × 963 = 621135 such pairs. We transform the drug-drug matrix into a drug-drugSE matrix. Thus, if the original drug-drug matrix has side effect *k* in the *ij*^th^ cell, then the transformed matrix has a 1 in the *i*^th^ row and (963(*j* – 1) + *k*)^th^ column. With this transformation, drugSE representations can be learnt through NCMF in addition to drug representations. We do not address the cold start problem and assume that all drug and drugSE instances in the test data (425883 in total) are present at least once in the train set. The memory on our system is insufficient to run NCMF directly on the transformed matrix collection. So, we first run just the ZINB autoencoder on the drug-drugSE matrix (of size 645 × 425883), on both the row and column dimensions, to get a lower-dimensional 32 × 1024 matrix. This matrix is combined with the drug-protein and protein-protein matrices and used as input to NCMF, to yield 50-dimensional drug representations. These drug representations and the 32-dimensional drugSE representations are used as features. For each triple (drug i – side effect k – drug j) in the training data, we use the representations of drug i and drugSE (j,k) with label 1 for positive triples and label 0 for negative triples, to train a Random Forest classifier. The trained classifier is used for predictions on the test data.

### 3 Experimental Results

#### 3.1 Runtime

The inference procedures in all the CMF methods are iterative by design. In the neural methods, DCMF and NCMF, training is done over multiple epochs. The number of iterations/epochs differ in each dataset depending on when the convergence criterion is met. The total time taken per dataset and the number of iterations/epochs are shown in Tables 12 and 13 respectively. The relative differences are seen in Table 11 itself, that shows the time taken per iteration or epoch. DFMF is the fastest; the neural methods NCMF and DCMF are comparable in speed and slower than DFMF. However, both DCMF and NCMF are faster than CMF and gCMF. Implementations of the methods are in R for CMF and gCMF, in python for DFMF and in pytorch (with GPU usage) for DCMF and NCMF. We note that a strict comparison would require ensuring uniformity in use of programming language and hardware. Nevertheless, this comparison provides a practical understanding of the runtimes involved with the available implementations.

**Table 11.** Time in seconds per iteration/epoch

| Dataset | NCMF | DFMF | DCMF | CMF | gCMF |
| --- | --- | --- | --- | --- | --- |
| PubMed | 38.2 | 1.1 | 34 | 501.9 | 947.8 |
| MIMIC | 2.4 | 0.1 | 3.6 | 39.1 | 38.6 |
| Polypharmacy | 0.7 | 0.01 | 0.3 | 2.8 | 2.9 |

**Table 12.** Time per dataset

| Dataset | NCMF | DFMF | DCMF | CMF | gCMF |
| --- | --- | --- | --- | --- | --- |
| PubMed | 13375 seconds | 108.762 seconds | 170 seconds | 1.162 days | 1.097 days |
| MIMIC | 2371 seconds | 9.048 seconds | 18 seconds | 2.175 hours | 1.072 hours |
| Polypharmacy | 354 seconds | 1.659 seconds | 1.4077 seconds | 9.508 mins | 4.99613 mins |

**Table 13.** Number of epochs

| Dataset | NCMF | DFMF | DCMF | CMF | gCMF |
| --- | --- | --- | --- | --- | --- |
| PubMed | 350 | 100 | 5 | 200 | 100 |
| MIMIC | 1000 | 100 | 5 | 200 | 100 |
| Polypharmacy | 490 | 100 | 5 | 200 | 100 |

**Table 14.** ARI for each of the 5 most frequent target proteins

| Protein ID | 1128 | 768 | 1269 | 1142 | 3269 |
| --- | --- | --- | --- | --- | --- |
| NCMF | -0.043795 | -0.001245 | 0.690685 | -0.002904 | 0.055541 |
| DCMF | -0.062869 | 0.008015 | 0.883528 | 0.000758 | 0.143492 |

### 3.2 Clustering

The ARI values in 5 for the PolyP2 dataset are averages obtained from the values in Table 3.2 for 5 most frequently associated target proteins in the data.

### 3.3 Polypharmacy Side Effect Prediction

Table 15 shows the top 10 best performing side effects and bottom 10 worst performing side effects with respect to the AUPRC scores achieved by NCMF. Such an analysis was also done for their model Decagon by Žitnik *et al*. (2018). Similar to their analysis we find that side effects in the bottom 10 are those with non-molecular origins and/or are common complaints, such as feeling unwell, headache and diarrhea. Their and our top 10 lists have several side effects in common, such as carbuncle, coccydynia and tympanic membrance perforation. A clear distinction between our score ranges and theirs can be observed: our worst performing scores range from 0.804 to 0.814 while theirs range from 0.679 to 0.712; similarly our best performing scores range from 0.996 to 0.999, while theirs range from 0.908 to 0.964. This is not surprising since both ESP and WDW were found to outperform Decagon on the same dataset and NCMF outperforms ESP and WDW.

**Table 15.** Best and worst performing side effects based on AUPRC.

| Worst Performing |  | Best Performing |  |
| --- | --- | --- | --- |
| Side Effect | AUPRC | Side Effect | AUPRC |
| NAUSEA | 0.804044 | SPLENECTOMY | 0.996205 |
| ASPARTATE_AMINOTRANSFERASE_INCREASE | 0.804452 | METHAEMOGLOBINAEMIA | 0.996333 |
| ARTERIAL_PRESSURE_NOS_DECREASED | 0.805014 | NASAL_POLYP | 0.996559 |
| ANAEMIA | 0.806016 | CARBUNCLE | 0.997082 |
| BODY_TEMPERATURE_INCREASED | 0.808163 | TYMPANIC_MEMBRANE_PERFORATION | 0.997171 |
| FEELING_UNWELL | 0.810073 | SKIN_STRIAE | 0.997221 |
| DIFFICULTY_BREATHING | 0.810612 | CUTANEOUS_CANDIDIASIS | 0.997649 |
| DIARRHEA | 0.812514 | PRIMARY_BILIARY_CIRRHOSIS | 0.998471 |
| HEAD_ACHE | 0.812977 | COCCYDYNIA | 0.999326 |
| DRUG_TOXICITY_NOS | 0.814319 | FASCIITIS | 0.999533 |

